# Persistence of SARS CoV-2 S1 Protein in CD16+ Monocytes in Post-Acute Sequelae of COVID-19 (PASC) Up to 15 Months Post-Infection

**DOI:** 10.1101/2021.06.25.449905

**Authors:** Bruce K. Patterson, Edgar B. Francisco, Ram Yogendra, Emily Long, Amruta Pise, Hallison Rodrigues, Eric Hall, Monica Herrara, Purvi Parikh, Jose Guevara-Coto, Timothy J. Triche, Paul Scott, Saboor Hekmati, Dennis Maglinte, Xaiolan Chang, Rodrigo A Mora-Rodríguez, Javier Mora

## Abstract

The recent COVID-19 pandemic is a treatment challenge in the acute infection stage but the recognition of chronic COVID-19 symptoms termed post-acute sequelae SARS-CoV-2 infection (PASC) may affect up to 30% of all infected individuals. The underlying mechanism and source of this distinct immunologic condition three months or more after initial infection remains elusive. Here, we investigated the presence of SARS-CoV-2 S1 protein in 46 individuals. We analyzed T-cell, B-cell, and monocytic subsets in both severe COVID-19 patients and in patients with post-acute sequelae of COVID-19 (PASC). The levels of both intermediate (CD14+, CD16+) and non-classical monocyte (CD14Lo, CD16+) were significantly elevated in PASC patients up to 15 months post-acute infection compared to healthy controls (P=0.002 and P=0.01, respectively). A statistically significant number of non-classical monocytes contained SARS-CoV-2 S1 protein in both severe (P=0.004) and PASC patients (P=0.02) out to 15 months post-infection. Non-classical monocytes were sorted from PASC patients using flow cytometric sorting and the SARS-CoV-2 S1 protein was confirmed by mass spectrometry. Cells from 4 out of 11 severe COVID-19 patients and 1 out of 26 PASC patients contained ddPCR+ peripheral blood mononuclear cells, however, only fragmented SARS-CoV-2 RNA was found in PASC patients. No full length sequences were identified, and no sequences that could account for the observed S1 protein were identified in any patient. Non-classical monocytes are capable of causing inflammation throughout the body in response to fractalkine/CX3CL1 and RANTES/CCR5.

## INTRODUCTION

Post-acute sequelae SARS-CoV-2 infection (PASC) is a disabling and sometimes debilitating condition that occurs in 10%-30% of individuals infected by SARS-CoV-2 and has recently been proposed to cause neurologic symptoms in 30% of those infected (*1*). The number and extent of symptoms is extremely heterogeneous with some reports suggesting >200 different symptoms (*2*). The underlying cause of PASC symptoms has remained a mystery though some data has pointed to tissue reservoirs of persistent SARS-CoV-2 as a potential mechanism (*3,4*). We recently reported a machine learning approach that identified the unique immunologic signature of individuals with PASC (*5*). In the same report, we also identified characteristic immune cell subset abnormalities that accompanied the unique cytokine/chemokine profile. The predominant immune cell abnormality was elevations in monocyte subsets. Monocyte subpopulations are divided into 3 phenotypic and functionally distinct types. Classical monocytes exhibit the CD14++, CD16-phenotype, intermediate monocytes exhibit a CD14+, CD16+ phenotype, and the non-classical monocytes express CD14lo, CD16+ (*6,7*). Further they express very different cell surface markers as previously described. In particular, classical monocytes express high levels of the ACE-2 receptor, the putative receptor for SARS-CoV-2 (*8*). Intermediate and non-classical monocytes express very little ACE-2 receptor. Similarly, classical monocytes express low levels of the chemokine receptors CX3R1 and CCR5. Intermediate monocytes express high levels of CCR5 while non-classical monocytes express high levels of CX3R1. Here, we report kinetic differences in the proportions of monocyte subsets in severe cases and PASC, as well as the presence of SARS-CoV-2 protein unaccompanied by corresponding viral RNA in CD14lo, CD16+ monocytes in PASC patients up to 15 months post-acute SARS-CoV-2 infection.

## RESULTS

Similar to other inflammatory and infectious conditions such as sepsis, lupus erythematosis, and rheumatoid arthritis among others (*9*), we detected statistically significant increases (P<0.002) of intermediate CD14+, CD16+ monocytes in individuals with PASC compared to healthy controls. In addition, CD14lo, CD16+ non-classical monocytes were also significantly elevated in PASC (P=0.01). Neither intermediate nor non-classical monocytes were elevated in severe COVID-19 (Figure 1).

**Figure 1.**
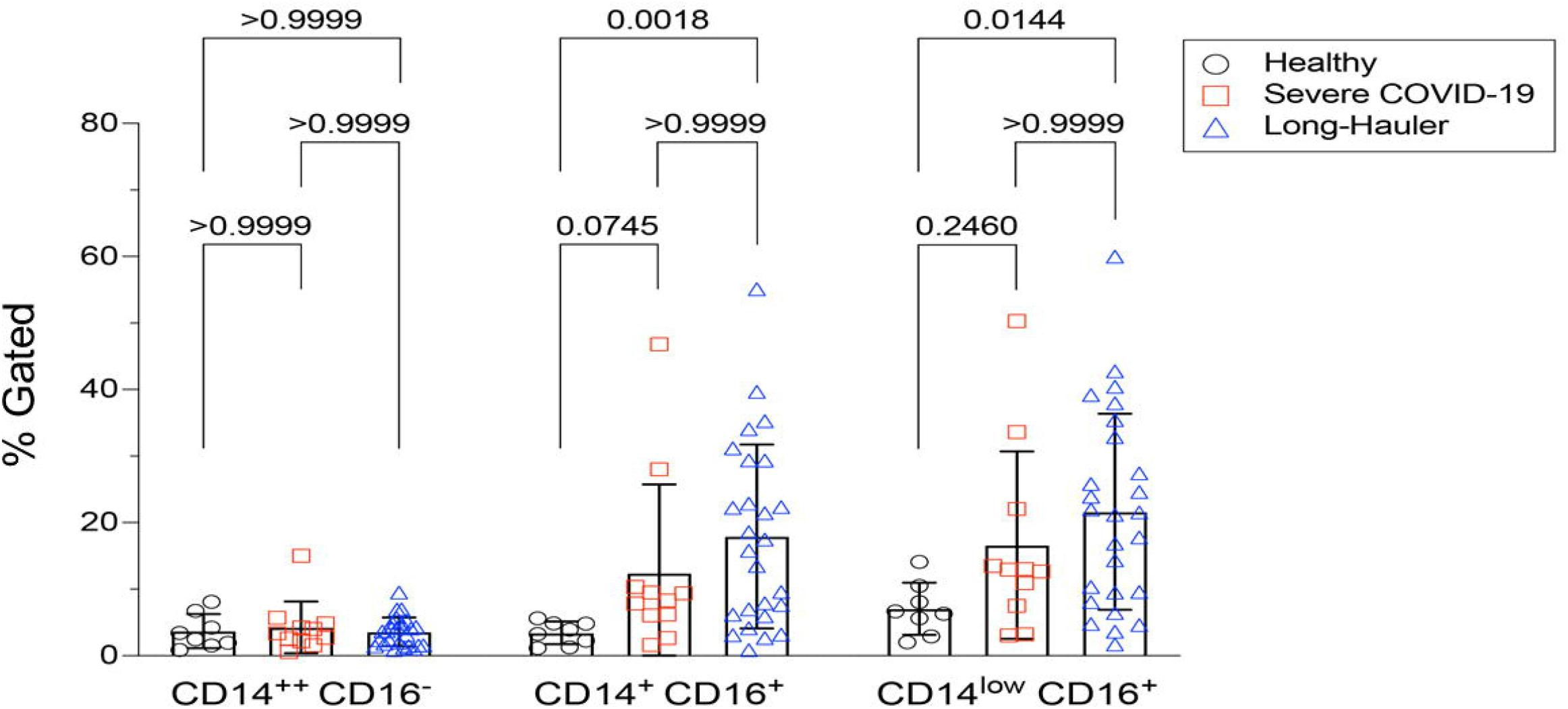
Quantification of classical, intermediate and non-classical monocytes in PASC (LH). Non-classical monocytes were significantly elevated in severe COVID-19 and in PASC.

Since the reports by our group and others found that monocyte subsets can be infected by HIV, HCV, Zika virus and Dengue fever virus (*10-12*), we screened peripheral blood mononuclear cells (PBMCs) from PASC individuals, as well as acute severe COVID-19 as controls, for SARS-CoV-2 RNA (Table 1). Using the highly sensitive, quantitative digital droplet PCR (ddPCR), we found that 36% (4 of 11) of severe COVID-19 patients’ PBMCs contained SARS-CoV-2 RNA compared to 4% (1/26) of PASC patients’ PBMCs. The one PASC patient that was RNA positive was 15 months post infection.

**Table 1.**
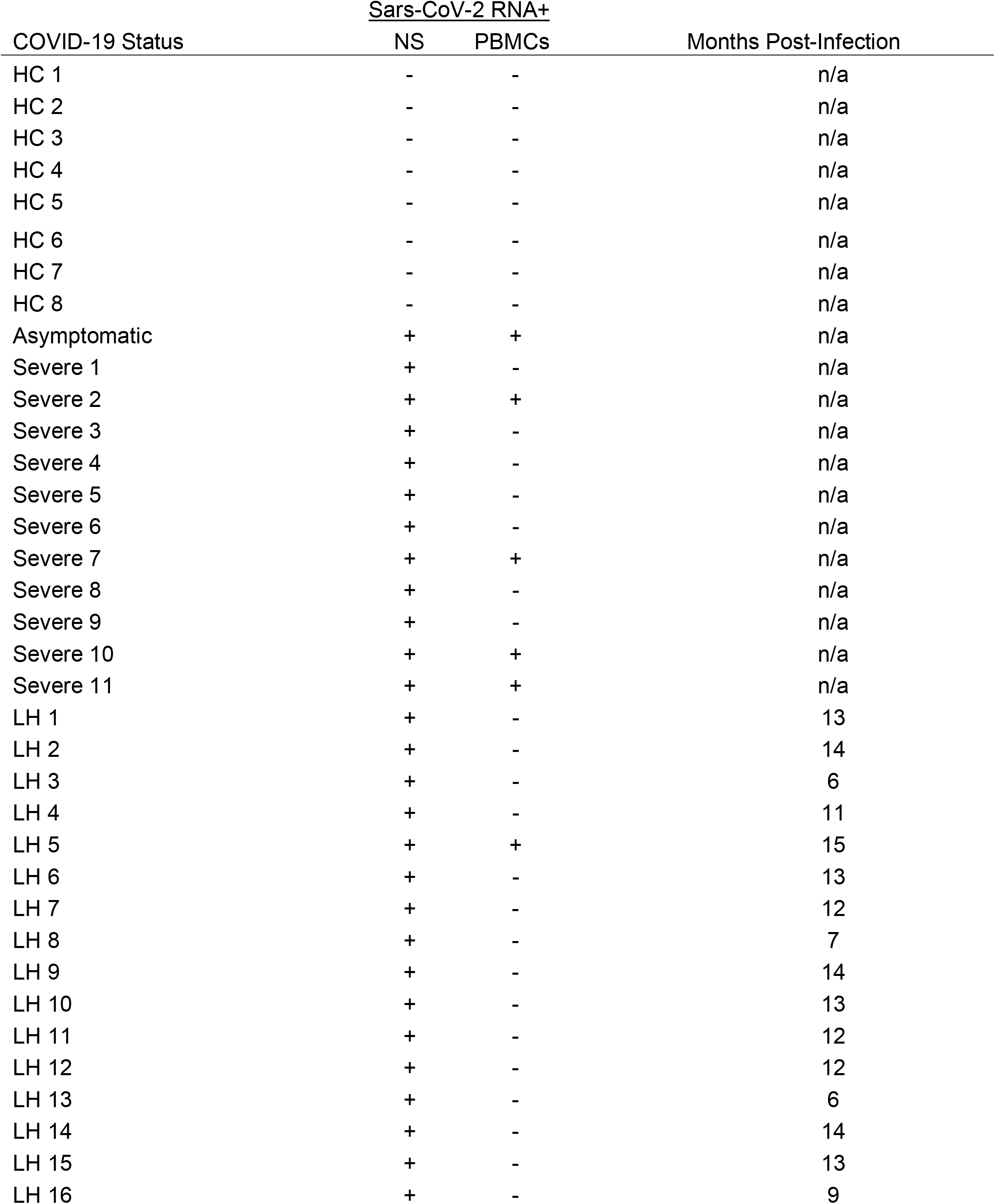
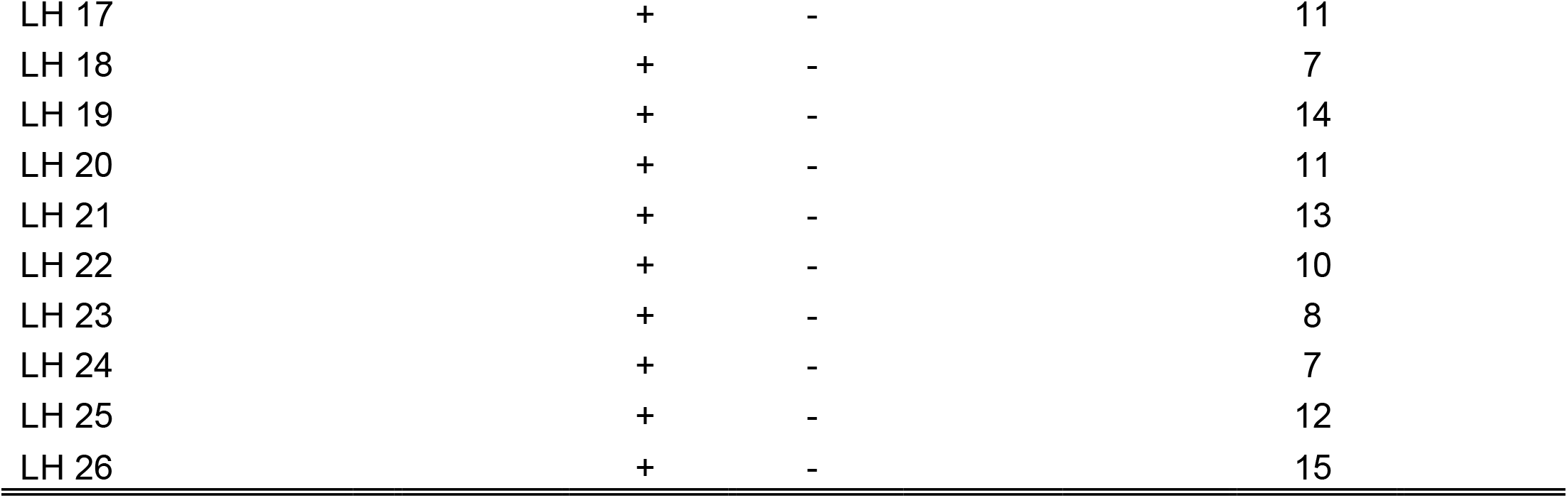
Molecular analysis of study participants.

To further establish the exact reservoir contributing to the positive signal detected using ddPCR, we performed high parameter flow cytometry with antibodies that define B cell, T-cell, and monocytic subsets in addition to simultaneous staining of these cells with an antibody for the SARS-CoV-2 S1 protein. As demonstrated in Figure 2, we found distinct subpopulations of SARS-CoV-2 containing cells in the CD14lo, CD16+ monocytic subset for 73% (19 out of 26) of PASC patients and 91% (10 out of 11) of severe COVID-19 patients. As demonstrated in Figure 3, the quantity of SARS-CoV-2 S1 containing cells were statistically significant in both the severe patients (P=0.004) and in the PASC patients (P=0.02). Neither classical monocytes nor intermediate monocytes expressed the SARS-CoV-2 S1 protein.

**Figure 2.**
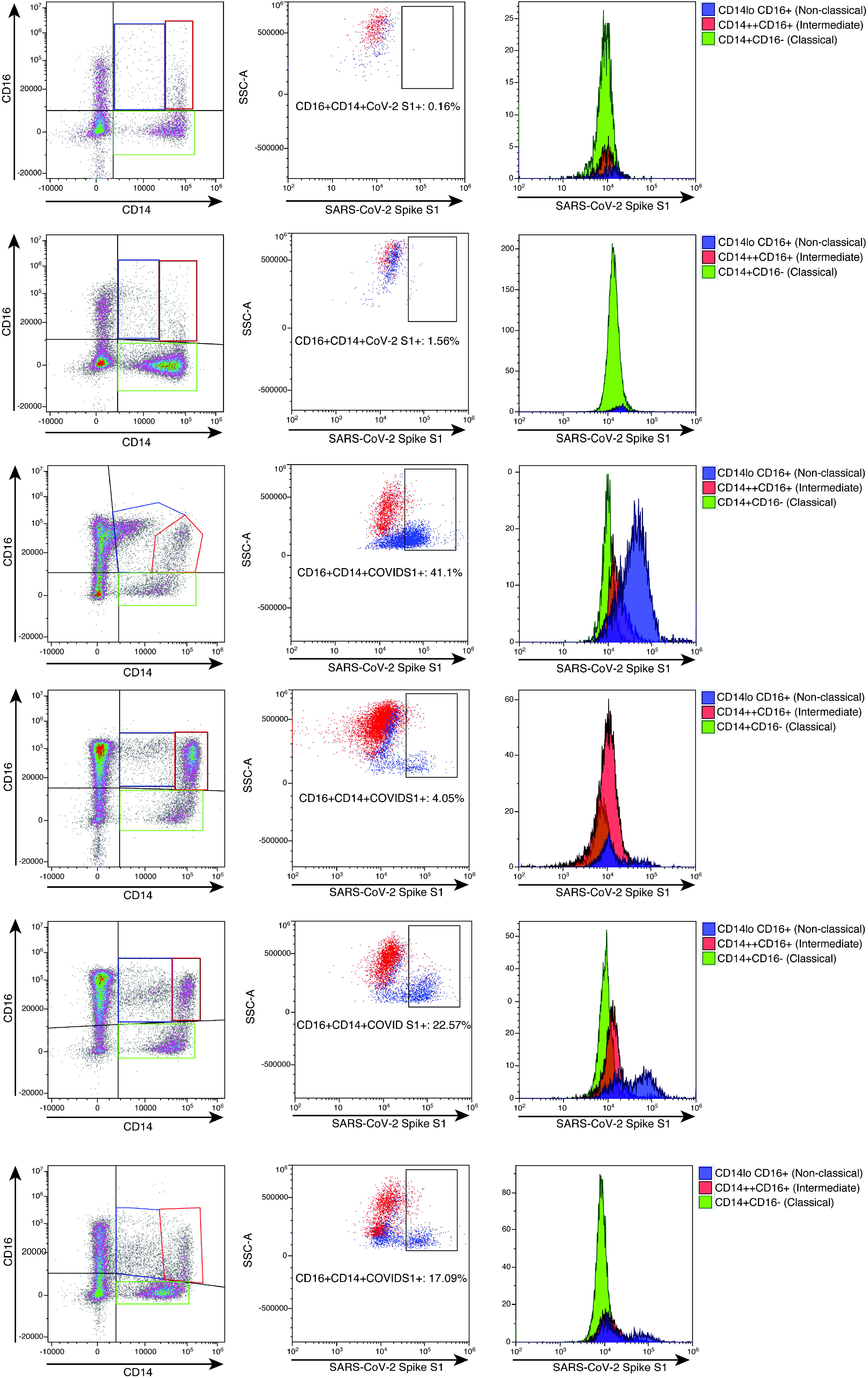
High parameter flow cytometric quantification of SARS-CoV-2 S1 protein in monocytic subsets. Cells were gated on CD45 then analyzed for CD14 and CD16 expression. Classical monocytes are green, intermediate monocytes are red and non-classical monocytes are blue.

**Figure 3.**
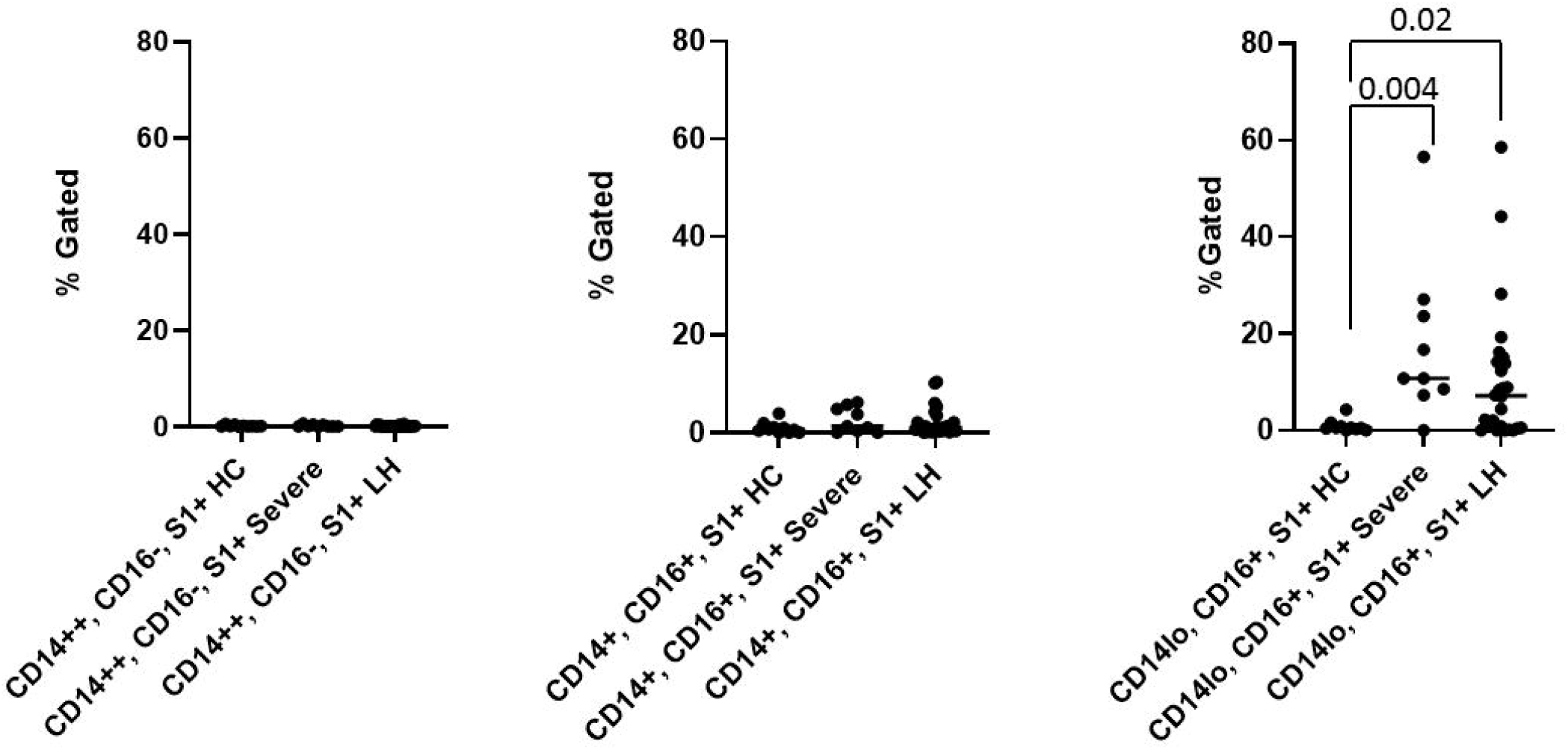
Quantification of SARS-CoV-2 S1 protein in monocyte subsets isolated from healthy controls (HC), severe COVID-19 (severe), and PASC patients (LH). SARS-CoV-2 S1 protein was expressed in non-classical monocytes in both severe and PASC individuals. The amount of expression was statistically significant.

To confirm the presence of SARS-CoV-2 S1 protein, we sorted CD14lo, CD16+ monocytes and performed Ultra High-Performance Liquid Chromatography (UHPLC). Following immunoprecipitation, the elution fractions were dried down *in vacuo*, resuspended in ddH_2_O and purified by to remove any non-crosslinked SARS-CoV-2 S1 antibody as well as any detergents from the commercial immunoprecipitation buffers. The UHPLC collected fractions were dried *in vacuo*, resuspended in 100 mM HEPES (pH 8.0, 20% Acetonitrile), and subjected to cistern: reduction and alkylation with chloroacetamide. The samples were then digested with AspN and LysC endopeptidases for 16h at 37°C. The digested peptides were analyzed on an Agilent 6550 IonFunnel QTOF and 1290 UHPLC by comparing patient samples to identical digests performed on commercially available SARS-CoV-2 S1 subunit. S1 subunit peptides from patient samples were mapped to a peptide database generated using commercial S1 subunit digests. Peptide identification consisted of matches in exact mass, isotope distribution, peptide charge state, and UHPLC retention time. As shown in Figure 4, the retention time of the representative peptide NLREFVFK in the digested commercial S1 subunit and Sample LH1-6 matched. Additionally, the Mass Spectra in Figure 4 show identical mass, isotope distribution, and charge states for the representative peptide NLREFVFK in the representative LH1 sample and commercial S1 subunit (also observed in LH 2-6, not shown). Using these metrics, up to 44% of the S1 subunit peptides could be identified in patient samples LH1-LH6 (Supplementary Table 1), providing complementary evidence to flow cytometry experiments that demonstrate the presence of S1 subunit protein in these patient cells.

**Figure 4.**
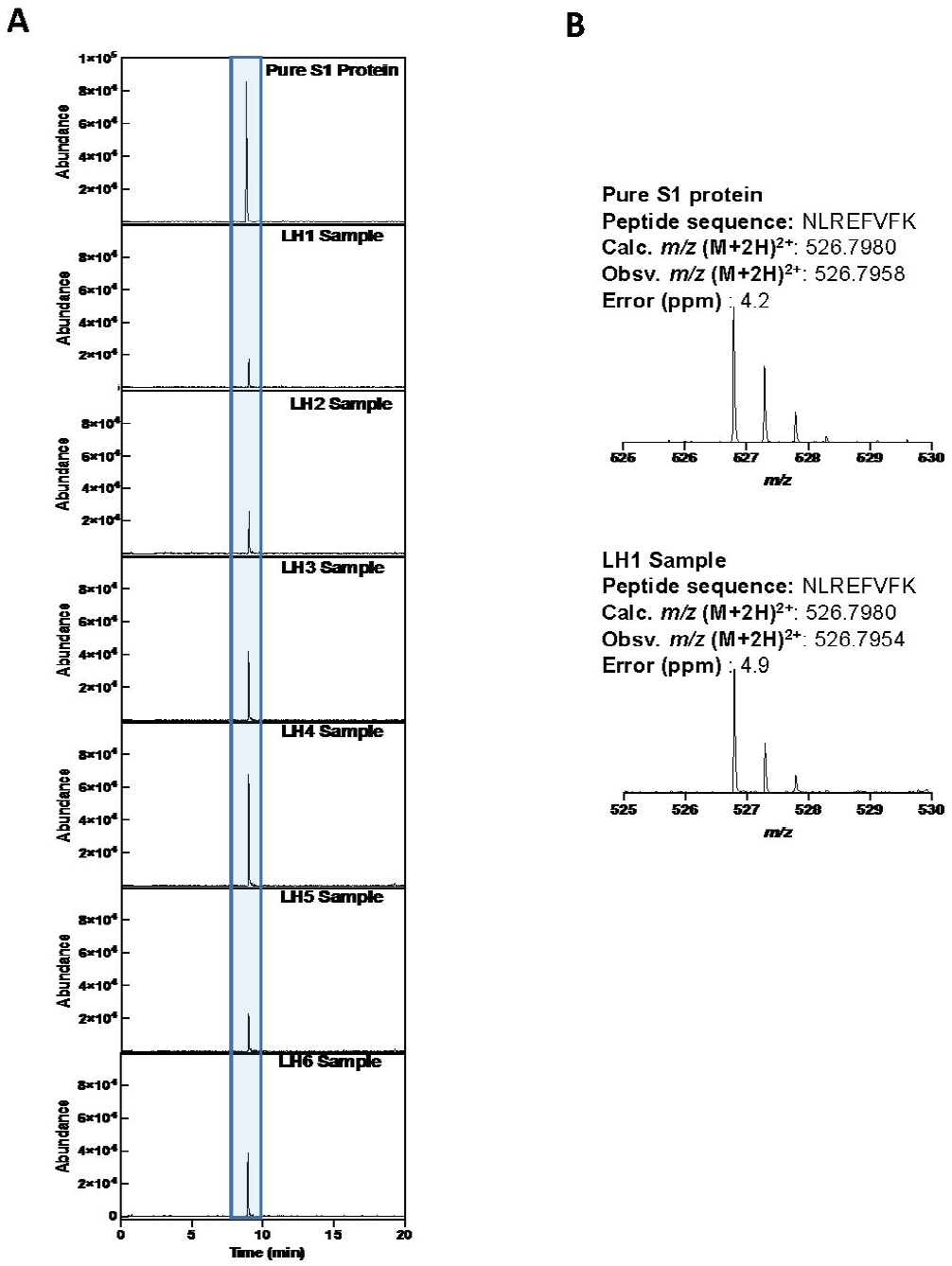
LCMS confirmation of the presence of S1 subunit in samples LH1-6. A. Extracted ion chromatogram (EIC) displaying the NLREFVFK peptide. The retention time matches that of the NLREFVFK peptide in the commercial S1 standard. B. Mass Spectra of the NLREFVFK from both the commercial standard and patient LH1. The Spectra show the same mass and isotope distribution.

To determine whether the observed S1 spike protein was a product of persistent viral infection, whole viral genome sequencing was performed on monocytes from five patients. Coverage analysis of the human control amplicons revealed adequate coverage to positively identify human genomic content. This is consistent with extraction of viral genomic content from a human host. Human controls also included targeted amplicons for amelogenin (*AMELX* and *AMELY*). The ratio of *AMELX* and *AMELY* reads is consistent with the known genders of each sample.

The sequencing coverage for the five samples was consistent with low viral titer samples or samples with high Ct values. Average coverage was between 24.17-592.87x and percent bases covered at 10x and 20x was between 10.81-19.18% and 7.69-15.24% respectively (Table 2). This is well below the expected threshold to eliminate stretches of Ns > 99 for consensus sequence submission to GenBank and > 90% genome coverage at 10x for accurate lineage determination and sequence submission to GISAID (www.gisaid.org). Evaluation of the reads revealed predominantly short reads (<100bp). To address poor quality reads, primer-dimers or reads that could possibly map to multiple loci, reads < MAPQ 10 were filtered resulting in the removal of 3.63-18.99% of total reads per sample.

**Table 2:** Average Coverage and Percent Bases Covered at 20x. While the percent of bases covered varied across patients, all were less than 20% at 10X, and less at 20X coverage. In no case was full length viral genome RNA detected, consistent with a lack of replication competent viral infection.

| Sample | Average Coverage | Percent Bases Covered at 10x | Percent Bases Covered at 20x |
| --- | --- | --- | --- |
| 02-03_20210625 | 171.64 | 19.18 | 15.24 |
| ABA-2_20210625 | 59.67 | 14.04 | 10.42 |
| BGI-2_20210625 | 24.17 | 10.81 | 7.69 |
| CST-2_20210625 | 40.29 | 11.71 | 7.79 |
| RG_20210625 | 592.87 | 12.6743 | 10.16 |

Lineage determination of the five samples from high quality mutations in the callable regions yielded lineages of *B* and *B*.*1* and were non-specific due to inadequate coverage across the genome. Mutations were identified in *ORF1ab* in all but sample LH5. LH5 had mutations in *N, S*, and *ORF3b*. (Figure 5).

**Figure 5:**
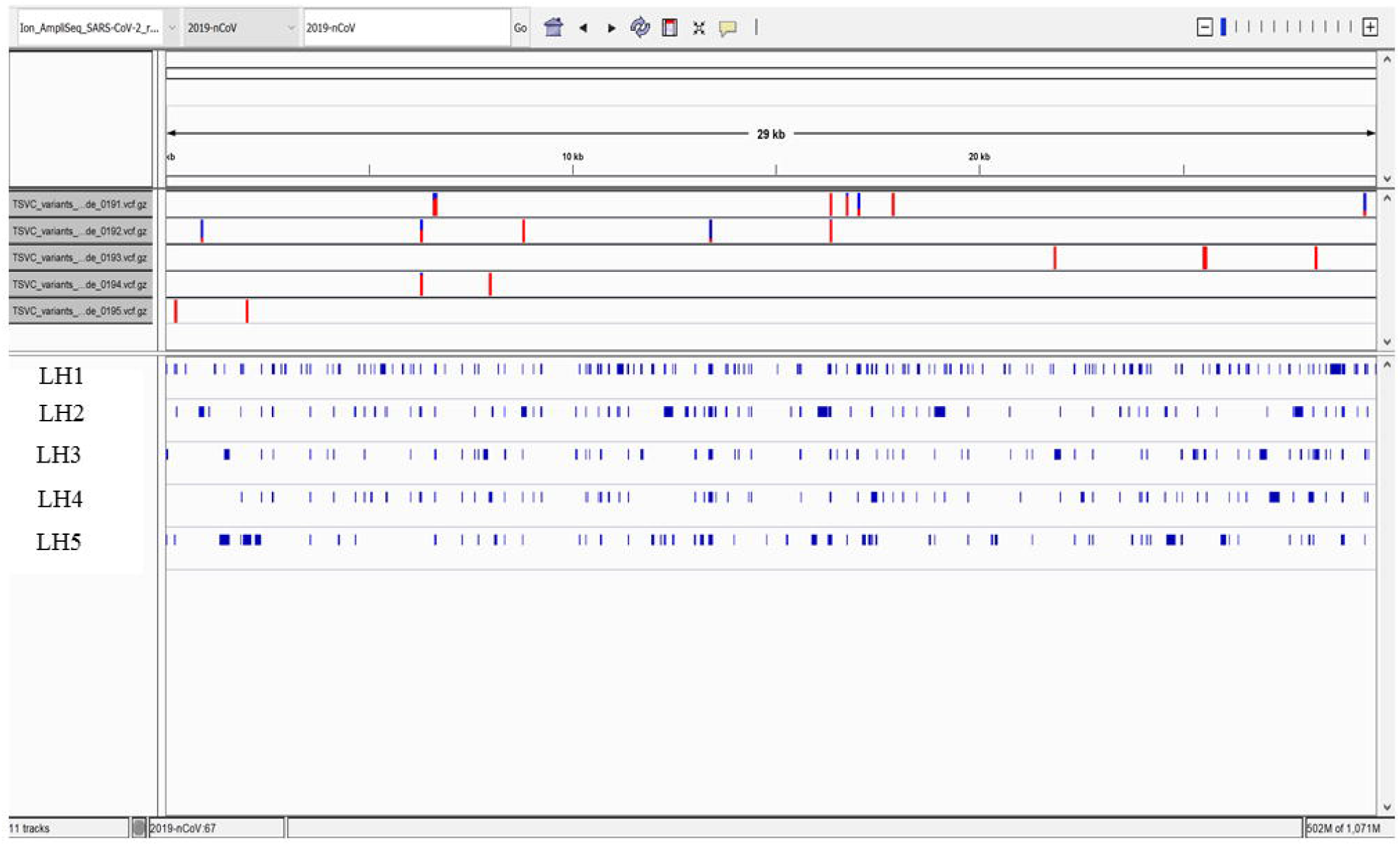
High Quality Mutations in the Callable Regions. Only fragmented viral RNA was identified in the five patients, but multiple mutations throughout the viral genome were identified, the vast majority of which were unique to each patient. Overall coverage was less than 20%, and no complete sequence in any portion of the viral genome was detected, including in the spike gene encoding the S1 subunit identified by protein analysis in these patients.

## DISCUSSION

Here, we report the discovery of persistent SARS-CoV-2 protein in CD14lo, CD16+ monocytes out to 15 months in some individuals and discuss the implications for the pathogenesis of PASC and severe cases of COVID-19. The three subtypes of circulating monocytes (classical, intermediate, non-classical) express very different cell surface molecules and serve very different functions in the immune system. Generally, classical’ monocytes exhibit phagocytic activity, produce higher levels of ROS and secrete proinflammatory molecules such as IL-6, IL-8, CCL2, CCL3 and CCL5. Intermediate monocytes express the highest levels of CCR5 and are characterized by their antigen presentation capabilities, as well as the secretion of TNF-α, IL-1β, IL-6, and CCL3 upon TLR stimulations. Non-classical monocytes expressing high levels of CX3CR1 are involved in complement and Fc gamma-mediated phagocytosis and anti-viral responses (*6*).

After maturation, human monocytes are released from bone marrow into the circulation as classical monocytes. Currently, strong evidence supports the concept that intermediate and non-classical monocytes emerge sequentially from the pool of classical monocytes (*13*). This is supported by transcriptome analysis showing that CD16+ monocytes have a more mature phenotype (*14*). In humans, 85% of the circulating monocyte pool are classical monocytes, whereas the remaining 15% consist of intermediate and nonclassical monocytes (*13*). Classical monocytes have a circulating lifespan of approximately one day before they either migrate into tissues, die, or turn into intermediate and subsequently nonclassical monocytes (*6,13*).

During pathologic conditions mediated by infectious/inflammatory reactions, the proportions of monocyte subsets vary according to the functionality of each specific subpopulation (*6,13,15*). Our previous results show that during early stages of the disease, PASC group have reduced classical monocyte and increased intermediate monocyte percentages compared with healthy controls (*5*). We find an increase in nonclassical monocytes in PASC group 6-15 months post infection, and higher percentages of intermediate and nonclassical monocytes at day 0 in severe cases, suggesting augmented classical-intermediate-nonclassical monocyte transition in both groups but with different kinetics.

The clinical relevance of monocyte activation in COVID-19 patients and the significance of these cells as viral protein reservoir in PASC is supported by our data reporting the presence of S1 protein within nonclassical monocytes. Viral particles and/or viral proteins can enter monocyte subpopulations in distinct ways, and this appears to be regulated differently in individuals that will develop severe disease or PASC. Classical monocytes are primarily phagocytes and express high levels of the ACE-2 receptor (*8*). Therefore, they could either phagocyte viral particles and apoptotic virally infected cells or be potential targets for SARS-CoV-2 infection. Considering their short circulating lifespan, viral protein-containing classic monocytes turn into intermediate and nonclassical monocytes. According to our results, this process happens faster in the severe group than in the PASC group. Indeed, at early stages of the disease the severe group show increased nonclassical monocytes whereas in PASC both the intermediate monocytes and non-classical monocytes are elevated. Additionally, CD14+CD16+ monocytes express intermediate levels of ACE-2 receptors and could as well serve as an infectious target of SARS-CoV-2 as it has been proved to be an infectious target of HIV-1 and HCV^11^. Nonclassical monocytes have been proposed to act as custodians of vasculature by patrolling endothelial cell integrity (*16*), thus pre-existing CD14lo CD16+ cells could ingest virally infected apoptotic endothelial cells augmenting the proportion of nonclassical monocytes containing S1 protein. This mechanism is more likely to take place in the PASC group where the S1 protein was detected 12-15 months post infection than in the severe group. Furthermore, nonclassical monocytes are associated with FcR-mediated phagocytosis (*17,18*), which might be related with the ingestion of opsonized viral particles after antibody production at later stages of the disease in PASC.

Previous reports indicate that the numbers of classical monocytes decrease, but the numbers of intermediate and non-classical monocytes increase in COVID-19 patients (*19*). Thus, the presence of S1 protein in nonclassical monocytes in both severe and PASC, might be associated with clinical characteristics and outcome of these groups. Previously, we found that individuals with severe COVID-19 have high systemic levels of IL-6, IL-10, VEGF and sCD40L (*5*). Consistent with our data, other studies showed association of increased production of IL-6, VEGF and IL-10 by nonclassical monocytes with disease severity (*20-22)*.

In the case of PASC, the persistence of circulating S1-containing nonclassical monocytes up to 15 months post infection, independently of the different possible mechanisms of viral proteins internalization discussed above, indicates that certain conditions are required to maintain this cell population. It has been shown in both humans and mice that nonclassical monocytes require fractalkine (CX3CL1) and TNF to inhibit apoptosis and promote cell survival (*22*). Our previous data show high IFN-γ levels in PASC individuals (*5*), which can induce TNF-α production (*23*). Further, TNF-α and IFN-γ induce CX3CL1/Fractalkine production by vascular endothelial cells^24^ creating the conditions to promote survival of nonclassical monocytes. Another important aspect is the permanency of S1-containing cells in the circulation, intermediate monocytes express high levels of CCR5 and extravasation of these cells can occur in response to CCL4 gradients. We showed that PASC individuals have low levels of CCL4 (*5*) maintaining these cells in circulation until they turn into nonclassical monocytes. Moreover, IFN-γ induced CX3CL1/Fractalkine production by endothelial cells (*23*) creates a gradient within the vascular compartment preserving nonclassical monocytes expressing CX3CR1 in the circulation.

Nonclassical monocytes are usually referred as anti-inflammatory cells (*22*), nevertheless it was recently shown that this subset can acquire a proinflammatory phenotype (*25*). Nonclassical monocytes acquire hallmarks of cellular senescence, which promote long term survival of these cells in circulation as explained above. Additionally, this induces an inflammatory state of the non-classical monocytes that could be a manifestation of the senescence-associated secretory phenotype (SASP), characterized by a high basal NF-κB activity and production of pro-inflammatory cytokines such as IL-1α, TNF-α and IL-8 (*25*).

The hallmark of PASC is the heterogeneity of symptoms arising in a variety of tissues and organs. These symptoms are likely associated with the inflammatory phenotype of these senescent nonclassical monocytes. The CD14lo, CD16+, S1 protein+ monocytes could be preferentially recruited into anatomic sites expressing fractalkine and contribute to vascular and tissue injury during pathological conditions in which this monocyte subset is expanded as previously demonstrated in non-classical monocytes without S1 protein. Previously, CD16+ monocytes were demonstrated to migrate into the brain of AIDS patients expressing high levels of CX3CL1 (fractalkine) and SDF-1 (*26*), and mediate blood-brain barrier damage and neuronal injury in HIV-associated dementia via their release of proinflammatory cytokines and neurotoxic factors. These sequelae are very common in PASC and these data could represent the underlying mechanism for the symptoms. Interestingly, a number of papers have been written discussing the increased mobilization of CD14lo, CD16+ monocytes with exercise (*27*). These data support the reports of worsening PASC symptoms in individuals resuming pre-COVID exercise regimens. In summary, the mechanism of PASC discussed in this report suggests that intermediate monocytes remain in circulation due to low CCL4 levels extending their time to differentiate leading to an accumulation of non-classical monocytes. The utility of using CCR5 antagonists in preventing migration of intermediate and non-classical monocytes due to the elevated levels of CCL5/RANTES in PASC (*5*). Further, our data suggests that interruption of the CX3CR1/fractalkine pathway would be a potential therapeutic target to reduce the survival of S1-containing non-classical monocytes and the associated vascular inflammation previously discussed (*5*) and presented here.

It is important to note that the S1 protein detected in these patients appears to be retained from prior infection or phagocytosis of infected cells undergoing apoptosis and is not the result of persistent viral replication. Full length sequencing of the five cases submitted for genomic analysis failed to identify any full-length sequence in the spike protein gene, or any other gene, that could account for the observed spike protein detected by proteomic analysis. In contrast, fragmented SARS-CoV-2 sequence was identified in all five of the cases. We have observed a pattern of high Ct value or negativity by PCR, accompanied by scant, fragmented viral sequence identified by whole viral genome sequencing over the past several months, a major shift from the low Ct value, full length viral sequences identified throughout most of 2020. The reasons for this shift are unclear, but as seen in these cases, it is unlikely these patients are producing any replication competent viral genomes, and are thus incapable of transmitting the infection. In contrast, the patients reported here appear to have developed an immune response to retained viral antigens, specifically the S1 fragment of the spike protein, which continues to be presented by CD16+ monocytes, eliciting an innate immune response characterized by elevated inflammatory markers including interferon ***γ***, IL-6, IL-10, and IL-2, among others. The body of evidence reported here would not support continued viral replication. Instead, it implicates dysregulation of innate immunity inflammatory mediators in response to persistent viral protein presentation by CD16+ monocytes.

## MATERIAL/METHODS

### Patients

Following informed consent, whole blood was collected in a 10 mL EDTA tube and a 10 mL plasma preparation tube (PPT). A total of 144 individuals were enrolled in the study consisting of 29 normal individuals, 26 mild-moderate COVID-19 patients, 25 severe COVID-19 patients and 64 chronic COVID (long hauler-LH) individuals. Long Haulers symptoms are listed in Figure 1. Study subjects were stratified according to the following criteria.

#### Mild

1. Fever, cough, sore throat, malaise, headache, myalgia, nausea, diarrhea, loss of taste and small
2. No sign of pneumonia on chest imaging (CXR or CT Chest)
3. No shortness of breath or dyspnea

#### Moderate

1. Radiological findings of pneumonia fever and respiratory symptoms
2. Saturation of oxygen (SpO2) ≥ 94% on room air at sea level

#### Severe

1. Saturation of oxygen (SpO2) < 94% on room air at sea level
2. Arterial partial pressure of oxygen (PaO2)/ fraction of inspired oxygen (FiO2) < 300mmHG
3. Lung infiltrate > 50% within 24 to 48 hours
4. HR ≥ 125 bpm
5. Respiratory rate ≥ 30 breaths per minute

#### Critical

1. Respiratory failure and requiring mechanical ventilation, ECMO, high-flow nasal cannula oxygen supplementation, noninvasive positive pressure ventilation (BiPAP, CPAP)
2. Septic Shock-Systolic blood pressure < 90mmHg or Diastolic blood pressure < 60 mmHg or requiring vasopressors (levophed, vasopressin, epinephrine
3. Multiple organ dysfunction (cardiac, hepatic, renal, CNS, thrombotic disease)

#### Post-acute COVID-19 (Long COVID)

1. Extending beyond 3 weeks from the initial onset of first symptoms

#### Chronic COVID-19

1. Extending beyond 12 weeks from the initial onset of first symptoms (Table 1S)

### High Parameter Immune Profiling/Flow Cytometry

Peripheral blood mononuclear cells were isolated from peripheral blood using Lymphoprep density gradient (STEMCELL Technologies, Vancouver, Canada). Aliquots 200 of cells were frozen in media that contained 90% fetal bovine serum (HyClone, Logan, UT) and 10% dimethyl sulfoxide (Sigma-Aldrich, St. Louis, MO) and stored at -70°C. Cells were stained and analyzed using a 17-color antibody cocktail including a PE-labeled SARS-CoV-2 S1 antibody (BioTechne, Minneapolis MN).

### Digital Droplet PCR

A QIAamp Viral Mini Kit (Qiagen, Catalog #52906) was used to extract nucleic acids from 300 to 400 mL of plasma sample according to the manufacturer’s instructions and eluted in 50 mL of AVE buffer (RNase-free water with 0.04% sodium azide). The purified nucleic acids were tested immediately with a Bio-Rad SARS-CoV-2 ddPCR Kit (Bio-Rad, Hercules, CA, USA). The panel was designed for specifically detecting 2019-nCoV (two primer/probe sets). An additional primer/probe set was used to detect the human RNase P gene in control samples and clinical specimens. RNA isolated and purified from the plasma samples (5.5 mL) was added to a master mix comprising 1.1 mL of 2019-nCoV triplex assay, 2.2 mL of reverse transcriptase, 5.5 mL of supermix, 1.1 mL of dithiothreitol, and 6.6 mL of nuclease-free water.

The mixtures were then fractionated into up to 20,000 nanoliter-sized droplets in the form of a water-in-oil emulsion in a QX200 Automated Droplet Generator (Bio-Rad, Hercules, CA). The 96-well real-time-digital droplet polymerase chain reaction (RT-ddPCR) ready plate containing droplets was sealed with foil using a plate sealer and thermocycled to reverse transcribe the RNA, before PCR amplification of cDNA in a C1000 Touch thermocycler (Bio-Rad, Hercules, CA, USA). After PCR, the plate was loaded into a QX200 Droplet Reader (Bio-Rad, Hercules, CA, USA) and the fluorescence intensity of each droplet was measured in two channels (FAM and HEX). The fluorescence data were then analyzed with QuantaSoft 1.7 and QuantaSoft Analysis Pro 1.0 Software (Bio-Rad, Hercules, CA, USA).

### Flow Cytometric Cell Sorting

Cryopreserved PBMCs were quick-thawed, centrifuged, and washed in 2% BSA solution in D-PBS. Cells were blocked for 5 min. in 2% BSA and then incubated at room temperature for 30 min. with Alexa Fluor® 488 Anti-CD45 antibody (IncellDx, 1/100 dilution), 2.5 ug of Alexa Fluor® 647 Anti-CD16 antibody (BD, Cat. # 55710), and 1 ug of PerCP/Cy5.5 Anti-human CD14 antibody (Biolegend, Cat. #325622). Cells were washed twice with 2% BSA/D-PBS, filtered, and kept on ice for the duration of the cell sort. Data was acquired on a Sony SH800, and only CD45+ cells staining positive for both CD14+ and CD16+ were sorted into test tubes with 100 uL 2% BSA solution. Sort purity of control PBMCs was confirmed to be >99% by re-analyzing sorted PBMCs using the same template and gating strategy.

### Single Cell Protein Identification

Patient cells were sorted based on phenotypic markers (as above) and frozen at -80° C. Six patient samples with positive flow cytometry signal and sufficient cell counts were chosen for LCMS confirmation. Frozen cells were lysed with the IP Lysis/Wash Buffer from the kit according to the manufacturer’s protocol. 10 ug of anti-S1 mAb were used to immunoprecipitate the S1 Spike protein from cell lysate of each patient. After overnight incubation with end-over-end rotation at 4°C and then three washes with IP Lysis/Wash Buffer, bound S1 Spike protein was eluted with the elution buffer from the kit.

IP elution fractions were dried *in vacuo*, resuspended in 20 uL of water, pooled, and purified by Agilent 1290 UPLC Infinity II on a Discovery C8 (3cm x 2.1 mm, 5 µm, Sigma-Aldrich, room temperature) using mobile phase solvents of 0.1% trifluoroacetic acid (TFA) in water or acetonitrile. The gradient is as follows: 5-75% acetonitrile (0.1% TFA) in 4.5 min (0.8 mL/min), with an initial hold at 5% acetonitrile (0.1% TFA) for 0.5 min (0.8 mL/min). The purified protein was dried *in vacuo* and resuspended in 50 µL of 100 mM HEPES, pH 8.0 (20% Acetonitrile). 1 µL of TCEP (100 mM) was added and the samples were incubated at 37°C for 30 min. 1 µL of chloroacetamide (500 mM) was added to the samples and incubated at room temperature for 30 min. 1 µL rAspN (Promega 0.5 µg/µL) and 1 µL of LysC (Pierce, 1 µg/µL) were added and the samples incubated at 37°C for 16 h, prior to LCMS analysis.

### LC-MS analysis

Digested recombinant SARS-CoV-2 Spike S1 protein was analyzed by a high mass accuracy mass spectrometer to generate a list of detectable peptides with retention time and accurate masses. An Agilent 1290 Infinity II high pressure liquid chromatography (HPLC) system and an AdvanceBio Peptide Mapping column (2.1 × 150 mm, 2.7 μm) were used for peptide separation prior to mass analysis. The mobile phase used for peptide separation consists of a solvent A (0.1% formic acid in H_2_O) and a solvent B (0.1% formic acid in 90% CH_3_CN). The gradient was as follows: 0–1 min, 3% B; 1– 30 min, to 40% B; 30–33 min, to 90% B; 33-35 min, 90% B; 37-39 min, 3% B. Eluted peptides were electrosprayed using a Dual JetStream ESI source coupled with the Agilent 6550 iFunnel time-of-flight MS analyzer. Data was acquired using the MS method in 2 GHz (extended dynamic range) mode over a mass/charge range of 50–1700 Daltons and an auto MS/MS method. Acquired data were saved in both centroid and profile mode using Agilent Masshunter Workstation B09 Data acquisition Software. The same analytical method was applied to immunoprecipitated samples from sorted patient cells except no ms/ms was acquired.

#### Viral Genome Detection by PCR and Whole Viral Genome Sequencing

##### Ct Determination with TaqPath Assay

Five RNA samples were subjected to the TaqPath COVID-19 Combo Kit Assay (Thermo Fisher Scientific Catalog no. A47814) to assess the cycle of threshold. TaqPath COVID-19 Combo Kit assay was performed according to recommendations of the EUA, using the Applied BioSystems QuantStudio 7 Flex (Thermo Fisher Scientific Catalog no. 4485701).

##### Whole Genome Sequencing of Samples with Ion AmpliSeq

Five RNA samples were subjected to AmpliSeq library preparation using the Ion AmpliSeq Library Kit 2.0 (Thermo Fisher Scientific Catalog no. 4480441) and the Thermo Fisher Scientific Insight panel, which consists of 238 amplicons in a two pool design against SARS-CoV-2 and seven amplicons as human controls. Libraries were prepared following manufacturer’s recommendations. Final libraries were amplified using 5 cycles of amplification and libraries were cleaned up using 0.5X right sided cleanup and 1.2X left sided cleanup using Kapa Pure Beads (Roche Catalog no l7983298001). Final libraries were quantified using Ion Library TaqMan Quantitation Kit (ThermoFisher Catalog no. 4468802). Samples were pooled in an equimolar distribution and loaded on to the Ion Chef Instrument (ThermoFisher Catalog no. 4484177) for Templating onto a 510 chip. The prepared chip was then loaded onto a GeneStudio S5 Prime (ThermoFisher Catalog no. A38196) for sequencing.

##### Genome Assembly, Quality Control, and Sequencing Analysis

Sequencing reads were aligned to the SARS-CoV-2 genome (build NC_045512.2) and human transcriptome (build GRCh37) using the Thermo Fisher Scientific TMAP aligner. Default parameters were used except for the *--context* flag.

Coverage analysis was performed by the coverage Analysis plugin in Thermo Fisher Scientific Torrent Suite software. Reads in the human controls were evaluated for quality control. Per-base coverage, average coverage, and percent genome covered at various depth thresholds were assessed using custom software. Read length distribution versus read quality (MAPQ score) were further evaluated.

Variant calling was performed on SARS-CoV-2 using the variantCaller plugin. Callable regions were identified as regions with read depth >= 20 after filtering reads with MAPQ < 10. Variants were filtered for quality by removing mutations with allele frequency (AF) < 0.5 in the callable regions. Lineage determination was made with pangoLEARN v1.2.13 using filtered-in mutations.

## Ethics

Informed consent was obtained from all participants.

## Data and materials availability

All requests for materials and raw data should be addressed to the corresponding author

## Author contributions

R.Y. and P.P. organized the clinical study and actively recruited patients.

B.K.P, A.P., H.R., X.E, E.L., J.B.S., TJT, PS, SH, DM performed experiments and analyzed the data.

J.G-C., R.A.M., J.M., X.C. performed the statistics and bioinformatics

B.K.P., J.M., EBF, J.G-C., R.A.M. wrote the draft of the manuscript and all authors contributed to revising the manuscript prior to submission.

## REFERENCES

1. R. Rubin. As Their Numbers Grow, COVID-19 “Long Haulers” Stump Experts. JAMA. 324, 1381–1383 (2020). doi:10.1001/jama.2020.17709

2. https://www.cdc.gov/coronavirus/2019-ncov/hcp/clinical-care/post-covid-conditions.html (April 8, 2021).

3. X.H. Yao, Z.C. He, T.Y. Li, H. R. Zhang, Y. Wang, H. Mou, Q. Guo, S.C. Yu, Y. Ding, X. Liu, Y.F. Ping, X.W. Bian. Pathological evidence for residual SARS-CoV-2 in pulmonary tissues of a ready-for-discharge patient. Cell Res. 30, 541–543 (2020). doi: 10.1038/s41422-020-0318-5. Epub 2020 Apr 28. PMID: 32346074; PMCID: PMC7186763.

4. R. Nienhold, Y. Ciani, V.H. Koelzer, A. Tzankov, J.D. Haslbauer, T. Menter, N. Schwab, M. Henkel, A. Frank, V. Zsikla, N. Willi, W. Kempf, T. Hoyler, M. Barbareschi, H. Moch, M. Tolnay, G. Cathomas, F. Demichelis, T. Junt, K.D. Mertz. Two distinct immunopathological profiles in autopsy lungs of COVID-19. Nat Commun. 11, 5086 (2020). doi: 10.1038/s41467-020-18854-2.

5. B.K. Patterson, J. Guevarra-Coto, R. Yogendra, E. B. Francisco, E. Long, A. Pise, H. Rodrigues, P. Parikh, J. Mora. R.A. Mora-Rodriguez. Immune-based prediction of COVID-19 severity and chronicity decoded using machine learning. Front Immunol (2021) https://doi.org/10.3389/fimmu.2021.700782.

6. T.S. Kapellos, L. Bonaguro, I. Gemünd, N. Reusch, A. Saglam, E. R Hinkley, J. L. Schultze. Human monocyte subsets and phenotypes in major chronic inflammatory diseases. Front Immunol 10, 1–13 (2020) https://doi.org/10.3389/fimmu.2019.02035.

7. L. Ziegler-Heitbrock. The CD14+ CD16+ blood monocytes: their role in infection and inflammation. J Leuk Biol 81, 584–592 (2007). https://doi.org/10.1189/jlb.0806510.

8. M. Rutkowska-Zapała, M. Suski, R. Szatanek, M. Lenart, K. Węglarczyk, R. Olszanecki, T. Grodzicki, M. Strach, J. Gąsowski, M. Siedlar. Human monocyte subsets exhibit divergent angiotensin I-converting activity Clin Exp Immunol 181, 126–132 (2015).

9. R. Mukherjee, P. Kanti Barman, P. Kumar Thatoi, R. Tripathy, B. Kumar Das, B. Ravindran. Non-Classical monocytes display inflammatory features: Validation in Sepsis and Systemic Lupus Erythematous. Sci Rep. 5, 13886 (2015). doi: 10.1038/srep13886.

10. D. Michlmayr, P. Andrade, K. Gonzalez, A. Balmaseda, E. Harris. CD14+CD16+ monocytes are the main target of Zika virus infection in peripheral blood mononuclear cells in a paediatric study in Nicaragua. Nature Microbiology, 2, 1462–1470 (2017). https://doi.org/10.1038/s41564-017-0035-0

11. Coquillard G, Patterson BK. Determination of hepatitis C virus-infected, monocyte lineage reservoirs in individuals with or without HIV coinfection. J Infect Dis. 200, 947–954 (2009). doi: 10.1086/605476. PMID: 19678757.

12. P. Ancuta, K.J. Kunstman, P. Autissier, T. Zaman, D. Stone, S.M. Wolinsky, D. Gabuzda. CD16+ monocytes exposed to HIV promote highly efficient viral replication upon differentiation into macrophages and interaction with T cells. Virology 344, 267–276 (2006). https://doi.org/10.1016/j.virol.2005.10.027

13. A.A. Patel, Y. Zhang, J.N. Fullerton, L. Boelen,, A. Rongvaux, A.A. Maini, V. Bigley, R.A. Flavell, D.W. Gilroy, B. Asquith, D. Macallan, S. Yona. The fate and lifespan of human monocyte subsets in steady state and systemic inflammation. J Exp Med. 214, 1913–1923 (2017). doi: 10.1084/jem.20170355. Epub 2017 Jun 12. PMID: 28606987; PMCID: PMC5502436.

14. P. Ancuta, K.Y. Liu Ky, V. Misra, V.S. Wacleche, A. Gosselin, X. Zhou, D. Gabuzda. Transcriptional profiling reveals developmental relationship and distinct biological functions of CD16+ and CD16-monocyte subsets. BMC Genomics 10, 403 (2009). doi: 10.1186/1471-2164-10-403. PMID: 19712453; PMCID: PMC2741492.

15. T. Tak, R. van Groenendael, P. Pickkers, L. Koenderman. Monocyte subsets are differentially lost from the circulation during acute inflammation induced by human experimental endotoxemia. J Innate Immun. 9, 464–474 (2017). doi: 10.1159/000475665. Epub 2017 Jun 23. PMID: 28641299; PMCID: PMC6738874.

16. C. Auffray, D. Fogg, M. Garfa, G. Elain, O. Join-Lambert, S. Kayal, S. Sarnacki, A. Cumano, G. Lauvau, F. Geissmann. Monitoring of blood vessels and tissues by a population of monocytes with patrolling behavior. Science 317, 666–670 (2007). doi: 10.1126/science.1142883. PMID: 17673663.

17. S.T. Gren, T.B. Rasmussen, S. Janciauskiene. A single-cell gene-expression profile reveals inter-cellular heterogeneity within human monocyte subsets. PLoS One 10, e0144351 (2015). doi: 10.1371/journal.pone.0144351. PMID: 26650546; PMCID: PMC4674153.

18. K.L. Wong, J.J. Tai, W.C. Wong, H. Han, X. Sem, W.H. Yeap, P. Kourilsky, S.C. Wong. Gene expression profiling reveals the defining features of the classical, intermediate, and nonclassical human monocyte subsets. Blood. 118, e16–31 (2011). doi: 10.1182/blood-2010-12-326355. Epub 2011 Jun 7. PMID: 21653326.

19. Jafarzadeh, P. Chauhan, B. Saha, S. Jafarzadeh, M. Nemati. Contribution of monocytes and macrophages to the local tissue inflammation and cytokine storm in COVID-19: Lessons from SARS and MERS, and potential therapeutic interventions. Life Sci. 257, 118102 (2020). doi: 10.1016/j.lfs.2020.118102. Epub 2020 Jul 18. PMID: 32687918; PMCID: PMC7367812.

20. Y. Zhou, B. Fu, X. Zheng. Pathogenic T-cells and inflammatory monocytes incite inflammatory storms in severe COVID-19 patients, National Science Review 7, 998–1002 (2020).

21. C.E. Olingy, C.L. San Emeterio, M.E. Ogle, J.R. Krieger, A.C. Bruce, D.D. Pfau, B.T. Jordan, S.M. Peirce, E.A. Botchwey. Non-classical monocytes are biased progenitors of wound healing macrophages during soft tissue injury. Sci Rep. 7, 447 (2017). doi: 10.1038/s41598-017-00477-1. PMID: 28348370; PMCID: PMC5428475.

22. P.B. Narasimhan, P. Marcovecchio, A.A.J. Anouk, C.C. Hedrick. Nonclassical monocytes in health and disease. Ann Rev Immunol 37, 439–456 (2019).

23. V. Vila-del Sol, C. Punzón, M. Fresno. IFN-gamma-induced TNF-alpha expression is regulated by interferon regulatory factors 1 and 8 in mouse macrophages. J Immunol 181:4461–4470 (2008). doi: 10.4049/jimmunol.181.7.4461. PMID: 18802049.

24. T. Matsumiya, K. Ota, T. Imaizumi, H. Yoshida, H. Kimura, K. Satoh. Characterization of Synergistic Induction of CX3CL1/Fractalkine by TNF-α and IFN-γ in vascular endothelial cells: an essential role for TNF-α in post-transcriptional regulation of CX3CL1. J Immunol 184, 4205–4214 (2010). DOI: 10.4049/jimmunol.0903212.

25. S.M. Ong, E. Hadadi, T.M. Dang, W.H. Yeap, C.T. Tan, T.P. Ng, A. Larbi, S.C. Wong. The pro-inflammatory phenotype of the human non-classical monocyte subset is attributed to senescence. Cell Death Dis 9, 266 (2008). https://doi.org/10.1038/s41419-018-0327-1.

26. C.F. Pereira, J. Middel, G. Jansen, H.S. Nottet. Enhanced expression of fractalkine in HIV-1 associated dementia. J Neuroimmunol 115, 168–115 (2001). doi: 10.1016/s0165-5728(01)00262-4. PMID: 11282167.

27. A.R. Jajtner, J.R. Townsend, K.S. Beyer, A.N. Varanoske, D.D. Church, L.P. Oliveira, K.A. Herrlinger, S. Radom-Aizik, D.H. Fukuda, J.R. Stout, J.R. Hoffman. Resistance exercise selectively mobilizes monocyte subsets: role of polyphenols. Med Sci Sports Exerc. 50, 2231–2241 (2018). doi: 10.1249/MSS.0000000000001703. PMID: 29957728.

